# *Cis*- and *trans*-eQTL TWAS of breast and ovarian cancer identify more than 100 risk associated genes in the BCAC and OCAC consortia

**DOI:** 10.1101/2023.11.09.566218

**Authors:** S. Taylor Head, Felipe Dezem, Andrei Todor, Jingjing Yang, Jasmine Plummer, Simon Gayther, Siddhartha Kar, Joellen Schildkraut, Michael P. Epstein

## Abstract

Transcriptome-wide association studies (TWAS) have investigated the role of genetically regulated transcriptional activity in the etiologies of breast and ovarian cancer. However, methods performed to date have only considered regulatory effects of risk associated SNPs thought to act in *cis* on a nearby target gene. With growing evidence for distal (*trans*) regulatory effects of variants on gene expression, we performed TWAS of breast and ovarian cancer using a Bayesian genome-wide TWAS method (BGW-TWAS) that considers effects of both *cis*- and *trans*-expression quantitative trait loci (eQTLs). We applied BGW-TWAS to whole genome and RNA sequencing data in breast and ovarian tissues from the Genotype-Tissue Expression project to train expression imputation models. We applied these models to large-scale GWAS summary statistic data from the Breast Cancer and Ovarian Cancer Association Consortia to identify genes associated with risk of overall breast cancer, non-mucinous epithelial ovarian cancer, and 10 cancer subtypes. We identified 101 genes significantly associated with risk with breast cancer phenotypes and 8 with ovarian phenotypes. These loci include established risk genes and several novel candidate risk loci, such as *ACAP3*, whose associations are predominantly driven by *trans*-eQTLs. We replicated several associations using summary statistics from an independent GWAS of these cancer phenotypes. We further used genotype and expression data in normal and tumor breast tissue from the Cancer Genome Atlas to examine the performance of our trained expression imputation models. This work represents a first look into the role of *trans-*eQTLs in the complex molecular mechanisms underlying these diseases.

## INTRODUCTION

Both breast and ovarian cancer carry a notable global burden. The estimated numbers of new cases of female breast cancer and ovarian cancer each year exceed 2.2 million and 310,000, respectively^1^. Genome-wide association studies (GWAS) have identified a growing catalog of validated common risk variants for breast and ovarian cancer^2–9^. Further research has helped define risk variants that are unique to distinct subtypes of breast cancer (for example, hormone receptor positive tumors) and ovarian cancer (for example, high grade and low grade serous histotypes)^5,6,8,10,11^. While pleiotropic and subtype-specific GWAS have helped delineate the germline genetic architecture of these cancers, most GWAS-derived risk variants for complex traits lie in non-coding regions of the genome^12,13^. This suggests that considerable disease risk may stem from variation in regulatory elements that affect gene transcription^14^.

Transcriptome-wide association studies (TWAS) are a powerful approach to identifying genes that are associated with risks for complex diseases with genetic effects mediated through genetically regulated transcriptional activity. In a training dataset, TWAS studies first build a statistical regression model of gene expression in a specific tissue by selecting those genetic variants having non-zero effect sizes on gene expression; we refer to such genetic variants as expression quantitative trait loci (eQTLs) of a broad sense for that gene. Using these models, TWAS then imputes the genetically regulated expression (GReX) levels of the gene in a target GWAS dataset where transcriptomic data are absent but disease outcome data are available. TWAS then tests for association between imputed gene expression and phenotype. Equivalent TWAS tests also can be conducted using only GWAS summary data with estimated eQTL effect sizes from the expression imputation models. TWAS have successfully identified novel candidate susceptibility genes for not only overall breast cancer and ovarian cancer risk, but, more recently, for specific subtypes of breast and ovarian cancer^15–17^.

To date, standard TWAS methods employ training models that only consider the regulatory effects of variants located in close proximity to the target gene (*cis*-SNPs)^18–24^. These variants reside within a small (e.g., 1Mb) window around the target gene. However, recent work has estimated the average proportion of heritability of gene expression estimated from these mapped *cis*-SNPs to be modest, with reported values ranging between 0.2 and 0.38^25^. One potential source for the remaining heritability of gene expression may be the aggregated effects of *trans*-eQTLs, which are defined as those variants that influence transcriptional activity that reside 1Mb or further away from the transcription start/end site of the target gene^25,26^. With growing evidence of distal regulatory effects of common variants, Luningham et al.^27^ developed a Bayesian genome-wide TWAS (BGW-TWAS) method, which trains expression prediction models considering both *cis*- and *trans*-SNPs. This approach both improves prediction of GReX as well as enhances detection of genes that influence phenotype through distal transcriptional regulation.

In light of established shared genetic etiology between breast cancer and ovarian cancer^13,28–32^, here we apply BGW-TWAS to conduct TWAS of each disease (and various subtypes) that consider the regulatory activity of both distal and proximal germline variants for a target gene. We first constructed gene expression imputation models using BGW-TWAS in normal breast and ovarian tissue from the Genotype-Tissue Expression (GTEx) project and subsequently imputed GReX into large-scale GWAS summary data from the Breast Cancer Association Consortium (BCAC) and Ovarian Cancer Association Consortium (OCAC) to identify genes associated with risk of overall breast cancer, five breast cancer subtypes (luminal A-like, luminal B-like, luminal B/HER2-negative-like, HER2-enriched-like, triple-negative), non-mucinous ovarian cancer, and five ovarian cancer subtypes (high grade serous, low grade serous, endometrioid, mucinous, clear cell). Our findings replicate several established cancer risk loci and suggest several novel candidate *trans*-eQTL driven genes not discovered by a standard TWAS approach that models *cis*-SNPs only. We then used independent GWAS summary data and matched genotype/gene expression data in breast tissue from the Cancer Genome Atlas to validate several of our top identified genes. This work provides new insight into the eQTL genetic architecture of breast and ovarian tissue and leverages *trans*-genome regulation of expression in these tissues for improved TWAS of breast and ovarian cancer.

## 2 MATERIALS AND METHODS

### 2.1 GTEx V8 training dataset

In order to train our imputation models for gene expression levels, we first obtained whole-genome sequencing (WGS) and RNA sequencing data on breast mammary tissue and ovarian tissue from the Genotype-Tissue Expression (GTEx) project V8 (dbGaP accession number phs000424.v8.p2). Data were available for 337 and 140 White individuals in breast tissue and ovarian tissue, respectively. We obtained gene expression levels as transcripts per million (TPM) per sample per tissue. We focused exclusively on autosomal genes for our analyses. We adjusted raw transcript data for the effects of age, body mass index, top five principal components, and top probabilistic estimation of expression residuals (PEER) factors. In the gene expression data from mammary tissue, we also adjusted for Estrogen Receptor 1 (*ESR1)* expression in accordance with previous studies^13,23^. This gene encodes estrogen receptor α, a transcription factor that plays a critical role in regulating gene expression and cell division in mammary glands^33^.

### 2.2 Breast cancer GWAS summary data

We obtained recently published summary-level GWAS data from BCAC (see Web Resources). The summary statistics for variant-level associations with breast cancer risk and risk of specific breast cancer subtypes were the result of a large multi-study GWAS of women of European ancestry^5^. The overall breast cancer analysis used genotype data from cases (invasive, *in situ*, unknown invasiveness) and controls across 82 BCAC studies that were genotyped using either the iCOGS or OncoArray Illumina genome-wide custom arrays. For this overall analysis, data from 11 other breast cancer GWAS were incorporated. This yielded a total sample size of 133,384 cases and 113,789 controls. The authors estimated SNP-disease associations using standard logistic regression, adjusting for country of origin and top principal components. Results were obtained for iCOGS subjects, OncoArray subjects, and additional GWAS separately and then combined via fixed-effects meta-analysis.

In addition to the summary statistics for the outcome of overall breast cancer risk, this study also published summary statistics for the association of variants with risk of specific intrinsic-like subtypes of breast cancer: luminal A-like cancer, luminal B-like cancer, luminal B/HER2-negative-like cancer, HER2-enriched-like, and triple-negative cancer. These subtype analyses were performed by fitting two-stage polytomous logistic regression models. Full details on these models are described elsewhere^34,35^. Only invasive cases were considered for this analysis, and samples from the 11 additional GWAS were not included due to missing tumor marker information. The final sample for the GWAS subtype analyses included 106,278 cases and 91,477 controls.

Lastly, a meta-analysis was performed combining results from the analysis of triple-negative breast cancer cases from BCAC and the separate analysis of cases and controls with a pathogenic *BRCA1* variant from the Consortium of Investigators of Modifiers of *BRCA1/2* (CIMBA). The CIMBA participants were also of European ancestry. Authors performed a fixed-effects meta-analysis, combining odds ratios estimates from the BCAC study and hazard ratios estimates from the CIMBA study. As the majority of breast cancer cases in *BRCA1* mutation carriers are triple-negative^36^, our “triple-negative” TWAS results utilized summary statistics from this meta-analysis for greatest power. Quality control and imputation protocols for all sets of genotype data used in the study are described separately^2,10,37,38^. Additional details on the study samples and statistical methods used in the breast cancer GWAS are provided by Zhang et al^5^.

Once we obtained the summary data from this GWAS, we performed liftover to map these GWAS variants to Human Genome Assembly GRCh38. We then harmonized and imputed missing variants’ summary statistics using the MetaXcan suite of tools (see Web Resources) and a European reference panel from the 1000 Genomes Project^39^.

### 2.3 Ovarian cancer GWAS summary data

We obtained summary-level GWAS data from a large-scale fine-mapping project of epithelial ovarian cancer^40^. This study used genotype data from six OCAC projects and two BCAC projects. The final sample contained a total of 26,151 ovarian cancer cases, 40,138 controls from OCAC, and 65,586 controls from BCAC. This project provided summary statistics for variant-level associations with risk of each of the five main subtypes of ovarian cancer: high grade serous ovarian cancer (HGSOC, 13,609 cases), low grade serous ovarian cancer (LGSOC, 2,749 cases), mucinous ovarian cancer (MOC, 2,587 cases), endometrioid ovarian cancer (ENOC, 2,877 cases), and clear cell ovarian cancer (CCOC, 1,247 cases). Additionally, the authors also performed a GWAS for the aggregate non-mucinous subtype, which excludes MOC cases. All participants were of European ancestry. Details on the genotyping and imputation procedures for this data are available elsewhere^40^.

Authors used logistic regression models to generate association statistics for SNP genotypes and the five subtype outcomes, as well as the non-mucinous epithelial ovarian cancer analysis. For each analysis, separate models were fitted for OncoArray data, COGS data, and five additional GWAS datasets that were also incorporated. Results from each group were then combined via fixed effects meta-analysis. Analyses adjusted for the effects of study of origin and possible population stratification by way of top principal components.

We performed liftover, harmonization, and summary statistic imputation as outlined for the breast cancer GWAS summary data.

### 2.4 Model training and association test

After we obtained the training dataset from GTEx V8 individuals with both WGS and RNA-seq transcriptomic data, we proceeded to train genome-wide expression prediction models separately in breast mammary tissue (N = 337) and ovarian tissue (N = 140) using BGW-TWAS. To briefly summarize, the method first calculates genome-wide single variant eQTL summary statistics from the simple linear regression of adjusted expression against genotype at a given SNP. *Cis*-SNPs were defined as those within 1Mb of the flanking 5’ and 3’ ends of the target gene. *Trans*-SNPs were those falling outside of the 1Mb window. Genotype files were then segmented into approximately independent genome blocks using LDetect^41^.

Next, we pruned genome-wide blocks to select a subset of blocks containing *cis-*SNPs and a subset of *trans* blocks with a minimum single-variant eQTL p-value less than 0.00001 to fit the following Bayesian variable selection regression (BVSR) model: ***E****_g_* = ***X****_cis_**W**_cis_* + ***X****_trans_**W**_trans_* + ε. Here, ***E****_g_* is vector of expression levels in the training (GTEx) dataset, ***X****_cis_* and ***X****_trans_* are the *cis* and *trans* genotype matrices from the pruned genotype blocks, and the w terms correspond to the eQTL effect sizes of the considered SNPs. The BGW-TWAS model then assumes a spike-and-slab prior distribution for the eQTL effect sizes, allowing these distributions to be different for *cis* and *trans* SNPs. Using an adapted expectation-maximization Markov Chain Monte Carlo algorithm, BGW-TWAS estimates (***w***, ***PP***), where, for selected SNPs, ***w*** is the vector of eQTL effects and ***PP*** is the vector of posterior causal probabilities (PP) of the selected SNPs being true eQTLs (with non-zero effect sizes). Only selected SNPs with estimated PP greater than 0.0001 were retained in the imputation models for each respective gene. Full details on the statistical methodology and computational algorithms of BGW-TWAS are provided by Luningham et al^27^.

Once we have trained the GReX imputation models, we then performed TWAS using the breast cancer and ovarian cancer GWAS summary statistics by calculating the following burden BGW-TWAS Z-score statistic for each gene *g*: 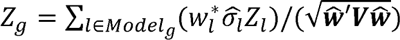. Here, 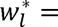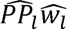, the expected eQTL effect size estimated from the BVSR model above. 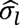 is a reference-derived estimate of standard deviation of genotype data for variant *l*, *Z_l_* is the variant’s corresponding Z-score statistic from the GWAS, and ***V*** is the reference-derived covariance of the genotype data of selected SNPs for this gene. We used GTEx V8 samples with available genotype information as reference for 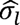 and ***V***. We calculated this burden BGW-TWAS Z-score statistic for each of the six breast cancer GWAS phenotypes and the six ovarian cancer GWAS phenotypes to test significant candidate risk genes.

Transcriptome-wide significant genes were those passing Bonferroni correction at 0.05/*M*. Here, *M* is the total number of gene-level association tests performed across all six analyses for breast cancer and ovarian cancer, respectively. We note that this is a strict multiple test correction, as we expect the gene-level test statistics to be correlated across certain breast cancer phenotypes and ovarian cancer phenotypes (such that the number of effectively independent tests is smaller than *M*). We further compared the performance of BGW-TWAS in identifying breast and ovarian cancer susceptibility genes to S-PrediXcan with pre-computed GTEx models that only consider *cis*-eQTLs^21^. We used the published GTEx V8 multivariate adaptive shrinkage in R (MASHR-M) models available at https://predictdb.org/. These models leverage information across multiple tissues and incorporate posterior causal probability of variants from fine-mapping. These models have been described in detail previously^42^. We applied S-PrediXcan to all 12 cancer phenotypes described above.

### 2.5 Validation analyses

#### 2.5.1 Independent genome-wide association dataset

To evaluate the robustness of our findings, we determined which genes identified by BGW-TWAS in the primary analyses using BCAC and OCAC GWAS data replicated in a similar analysis using summary statistic data from an independent GWAS of multiple cancers^43^. This GWAS aimed to identify common germline genetic variants associated with 18 types of cancer and interrogate possible pleiotropy among these identified variants. The study sample comprised of individuals of European ancestry from the UK Biobank (UKB) and the Kaiser Permanente Genetic Epidemiology Research on Adult Health and Aging (GERA) cohort. A total of 17,881 breast cancer cases (13,903 UKB, 3,978 GERA) and 1,259 ovarian cancer cases (1,006 UKB, 253 GERA) were considered, as well as a total of 219,656 controls (189,855 UKB, 29,801 GERA). Following quality control, imputation was performed using 1000 Genomes reference data. Authors fitted cohort-specific logistic regression models. These models adjusted for the effects of age, top 10 PCs, and genotyping array and genotype reagent kit (where applicable). Results were then combined via meta-analysis for variants present in both cohorts. We obtained the publicly available summary statistics for the fixed-effect meta-analysis of breast and ovarian cancer (see Web Resources). Again, we performed variant harmonization and liftover to GRCh38 using the MetaXcan suite of tools. We repeated BGW-TWAS with this data for significant genes identified in the primary BCAC/OCAC analyses. We note that this validation GWAS data reflects odds ratio estimates for overall breast cancer and ovarian cancer, as the GWAS study did not include subtype-specific regression models.

#### 2.5.2 GReX prediction in independent breast tissue samples

As a second validation analysis, we investigated how the GReX imputation models trained by BGW-TWAS in GTEx data (normal breast tissue) would predict gene expression in an independent set of breast cancer cases. Specifically, we looked at prediction performance in tumor-adjacent normal breast tissue (NAT) and tumor tissue samples. We obtained individual-level germline genotype data and matched gene expression data in breast cancer cases from The Cancer Genome Atlas (TCGA). Genotypes were called from Affymetrix SNP Array 6.0. We restricted consideration to individuals of consensus European ancestry as defined by Carrot-Zhang et al^44^. In this paper, authors performed four methods of ancestry determination in TCGA individuals. Consensus ancestry refers to the majority ancestry assignment across the employed methods. We also retrieved ancestral PCs for TCGA samples from this study. To maximize sample size in TCGA, we downloaded both blood-derived genotype data and solid normal tissue-derived (adjacent to tumor) genotype data. For individuals who had both, we preferentially used blood-derived genotypes as germline. We set as missing genotypes with birdseed confidence scores exceeding 0.1. We retained SNPs with call rate greater than 95% and samples with call rate greater than 95%. We aligned alleles to agree with Affymetrix SNP Array 6.0 annotation. We then excluded ambiguous SNPs and duplicates with identical chromosome and position and removed samples with high heterozygosity. We defined this when the absolute value of the inbreeding coefficient exceeded 0.2. We also pruned sample pairs with high estimated relatedness. We defined this by an estimated KING kinship coefficient exceeding 0.0884 (2^nd^ degree relatedness or higher)^45^. We excluded SNPs with MAF < 0.005 and HWE p < 1e-6. We aligned alleles with data from the 1000 Genomes. We performed imputation using the Michigan Imputation Server with 1000 Genomes Phase 3 V5 data and applied a Rsq threshold of 0.3. We then lifted variants over to GRCh38.

We downloaded RNA sequencing data (Illumina TruSeq) in NAT breast tissue and tumor tissue from TCGA. We ultimately had 786 individuals with germline genotype data, complete covariate information, and gene expression data in breast tumor samples. We had 101 individuals with complete genotype data, covariate data, and expression data in NAT breast tissue samples. We selected genes as having TPM ≥ 0.1 in 20% more of samples and those having 6 or more reads in 20% or more of samples. To calculate PEER factors, we then normalized read counts using the trimmed means of M values (TMM) method^46^ and applied inverse normal transformation. These factors were calculated using *peer* package in R^47^. For both models (NAT and tumor), we determined the number of PEER factors to adjust for by examining the plot of the posterior variance of factor weights for a clear elbow. Expression levels in tumor were adjusted for the effects of age, first 10 PCs, first 5 PEER factors, and *ESR1* expression via linear regression. For NAT samples, we adjusted for age, first 3 PCs, first 3 PEER factors, and *ESR1* expression.

Once all data were processed, we used our GTEx-derived *cis*- and *trans*-eQTL gene expression imputation models constructed for the main analysis to predict gene expression using TCGA germline variants. For genes that we had successfully trained a model for in Section 2.4 and were present in the TCGA expression datasets, we estimated the correlation between predicted GReX and observed, adjusted expression levels in both NAT and tumor samples. For this, we used Spearman’s rank correlation coefficient.

## 3 RESULTS

### 3.1 Fitted GReX models and eQTL architecture

In normal GTEx breast tissue samples, we successfully trained 24,833 autosomal gene expression models using Bayesian variable selection regression (BVSR) within BGW-TWAS. For each variant included in the BGW-TWAS models, we estimated the posterior probability (PP) of the variant being a true eQTL for the target gene. The sum of these PP estimates across all SNPs included as predictors in the fitted model corresponds to the estimated number of eQTLs per gene. These quantities can help improve our understanding of the location and distribution of eQTL effects genome-wide. For quality control, we excluded 102 outlier gene models with estimated PP summation exceeding 35 and/or maximum 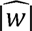 (eQTL effect size) greater than 1000, indicating poor model fit. Across the remaining genes passing these quality control filters, the median number of SNPs with estimated PP > 0.0001 per model was 868 (interquartile range [IQR] = 1074). For all breast tissue models, the average training R^2^ (squared correlation between imputed GReX and observed gene expression in the training GTEx dataset) was 0.30 (SD = 0.11). The median proportion of variants included as predictors per model that was located in *trans* to the target gene was 0.98 (IQR = 0.04). The median number of estimated eQTLs per gene was 1.14 (IQR = 2.04), while the median estimated eQTLs from *trans* regions across all breast models was 0.74 (IQR = 1.70). The distribution of estimated total eQTLs in breast tissue expression models is shown in Supplemental Figure 1. The median (IQR) of estimated eQTLs located in *trans* regions on the same chromosome as the target gene was 0.02 (0.05). The median (IQR) of estimated eQTLs located in *trans* regions on a different chromosome from the target gene was 0.69 (1.60). This distribution indicates most SNPs used to model gene expression in the *trans* genome were located on a different chromosome than the target gene. The median genome-wide, *cis*-region, and *trans*-region estimates of total eQTLs according to model training R^2^ are provided in Supplemental Table 1.

In normal ovarian tissue samples from GTEx, we trained 22,584 autosomal GReX models using the BVSR framework of BGW-TWAS. All models had cumulative PP below 35, but 68 models with large maximum 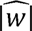 were excluded. Across our fitted BGW-TWAS models, the number of genome-wide variants included as predictor variables was generally smaller than for the breast tissue models, with a median of 608 (IQR = 983). The median proportion of variants included as predictors per model that located in *trans* to the target gene was 0.98 (IQR = 0.03). The median number of estimated eQTLs per gene was lower for genes in ovarian tissue at 0.40 (IQR = 1.27), while the median estimated eQTLs from *trans* regions across models was 0.30 (IQR = 1.14). The distribution of estimated total eQTLs in ovarian tissue expression models is also provided in Supplemental Figure 2. The median (IQR) of estimated eQTLs located in *trans* regions on the same chromosome as the target gene was 0.01 (0.04). The corresponding median (IQR) of estimated eQTLs residing on a different chromosome as the target gene was 0.28 (1.06). The median genome-wide, *cis*-region, and *trans*-region estimates of total eQTLs according to model training R^2^ are similarly provided in Supplemental Table 1. The average training R^2^ for fitted ovarian tissue models was 0.49 (SD = 0.13). We note that a larger proportion of genes had training R^2^ exceeding 0.5 in ovarian tissue compared to breast tissue. This inflation is likely a result of limited ovarian tissue samples with RNA sequencing data for model training, as observed in previous simulations for this method^27^.

### 3.2 Breast cancer TWAS

We performed a total of 148,929 tests for BGW-TWAS across the six breast cancer phenotypes. This corresponds to a Bonferroni-adjusted p-value threshold of 3.36×10^−7^ for transcriptome-wide significance. We note again this correction is stringent as all tests are not expected to be independent across phenotypes. We further obtained S-PrediXcan results based on *cis*-eQTLs only using the GTEx V8-derived MASHR models for 14,145 genes for each of the breast cancer phenotypes. We performed a total of 84,870 tests with S-PrediXcan, corresponding to a Bonferroni threshold of 5.90×10^−7^. Manhattan plots and quantile-quantile (QQ) plots of the BGW-TWAS results for the analysis of overall breast cancer risk and risk of the five common subtypes of breast cancer are included in Supplemental Figures 3-9. We see that the BGW-TWAS p-values do appear to suffer from inflation for the overall and luminal A phenotypes. As an illustrative example, for overall breast cancer, we have a genomic inflation factor of λ = 1.34, likely the result of the large GWAS sample sizes and phenotype polygenicity. Thus, as an alternative measure, we calculated the genomic inflation factor scaled to a study of 1000 cases and 1000 controls (λ_1000_ = 1.003)^48–50^. This measure was acceptable and reflective of all our BGW-TWAS results for breast cancer phenotypes.

Across the six TWAS performed for breast cancer phenotypes, BGW-TWAS identified 101 unique genes significantly associated with at least one of the six phenotypes considered. Location and phenotypes associated with these genes are shown in Figure 1. This figure also illustrates how the location of these genes relate to the location of significant GWAS variants from the original BCAC analysis of overall breast cancer risk. We then performed a validation TWAS of these 101 genes using independent GWAS summary statistics of breast cancer from Rashkin et al^43^. For overall breast cancer risk, we identified 87 significant genes initially from BCAC (see full list in Supplemental Table 2) and, of these, 31 further validated in the BGW-TWAS of Rashkin et al. data (p < 0.05/101). These 31 genes are provided in Table 1 along with sPrediXcan p-values. Of the 31 genes, 23 either could not be fit using MASHR-M models with sPrediXcan or failed to reach statistical significance in the *cis*-eQTL only approach.

**Figure 1.**
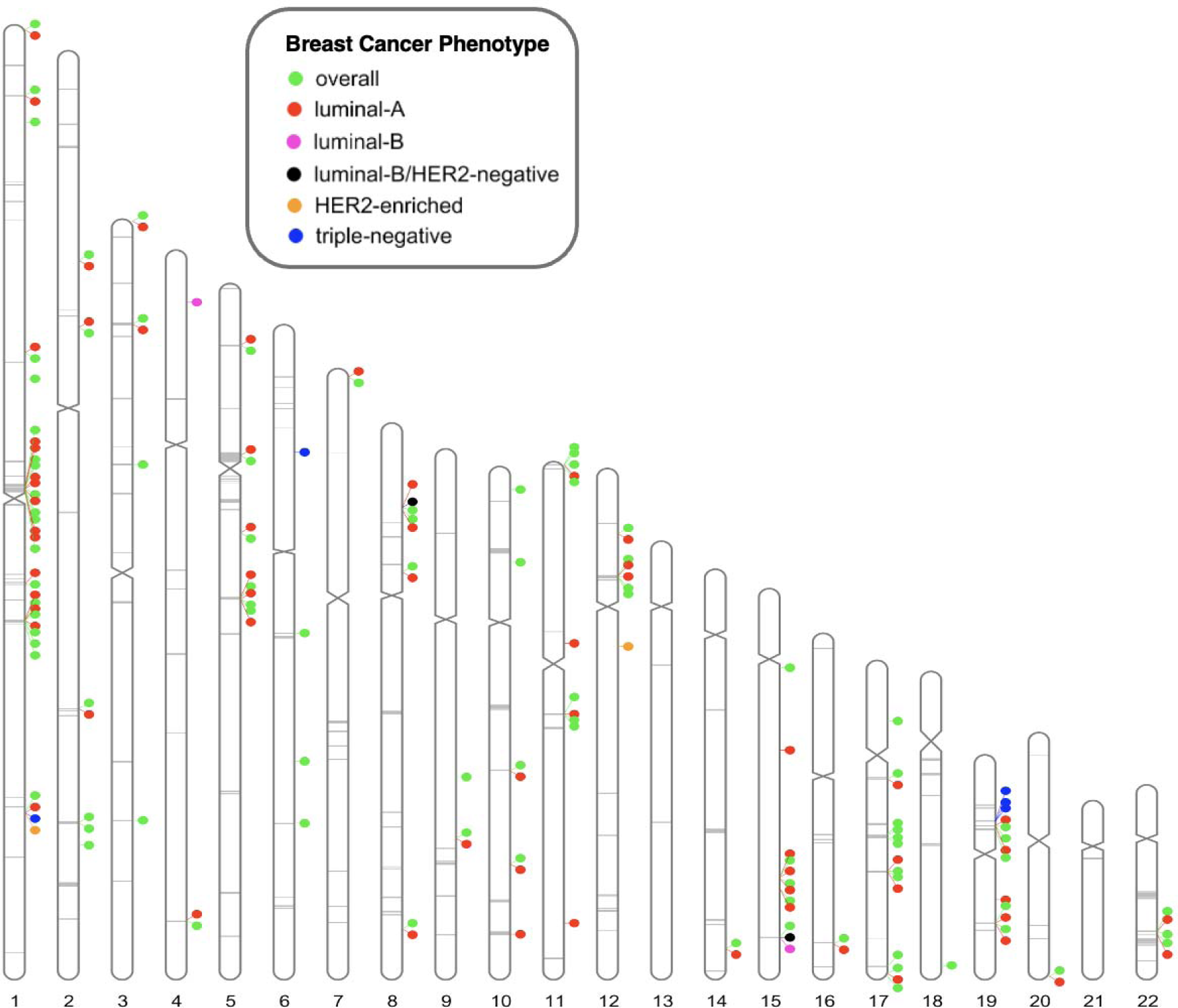
Ideogram of BCAC-derived BGW-TWAS results for overall breast cancer and breast cancer subtypes. 101 genes shown meet transcriptome-wide Bonferroni-adjusted p-value threshold for one or more phenotypes. Gray lines indicate position of genetic variants with BCAC GWAS p < 51710-8 for association with overall breast cancer risk.

**Table 1.**
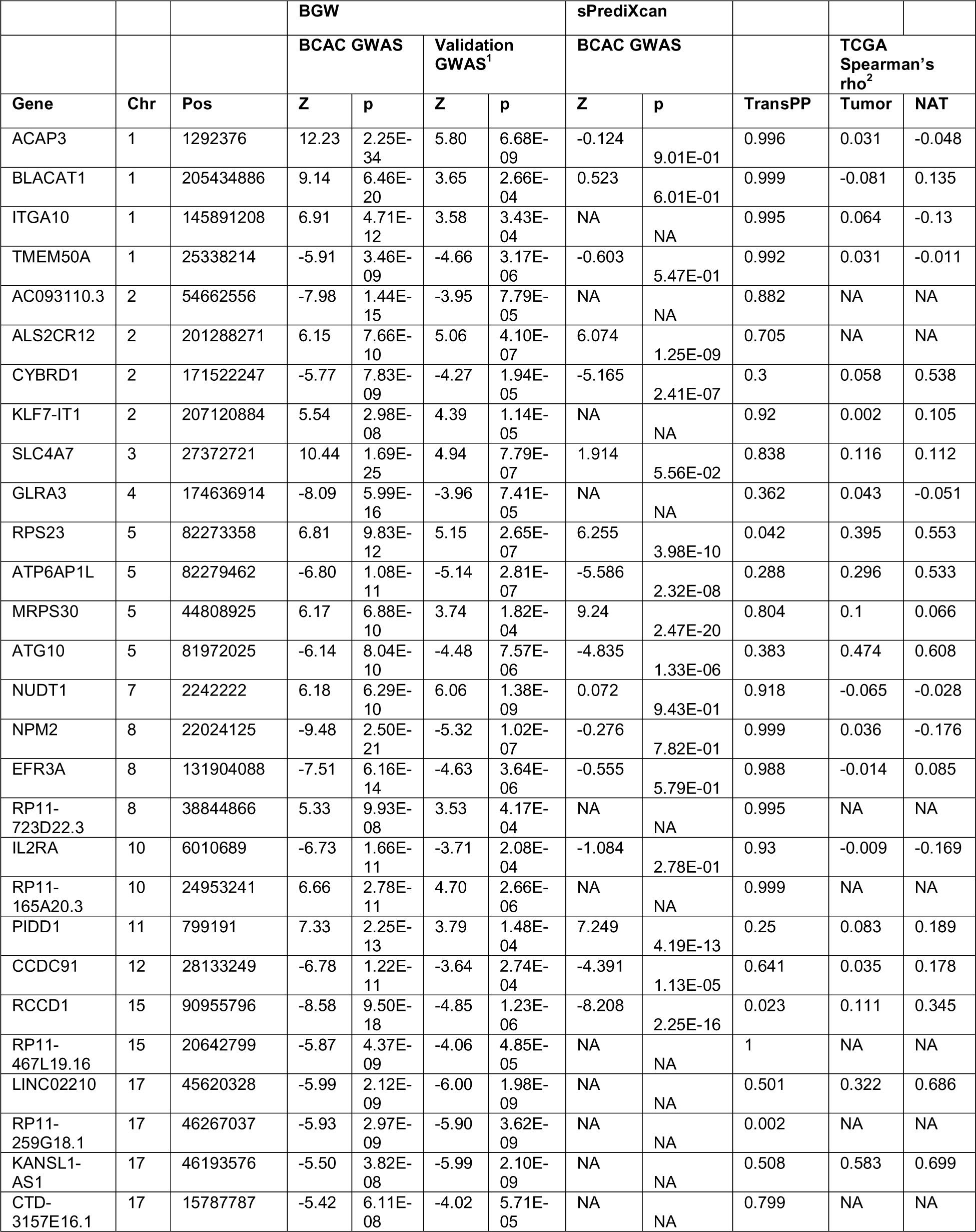
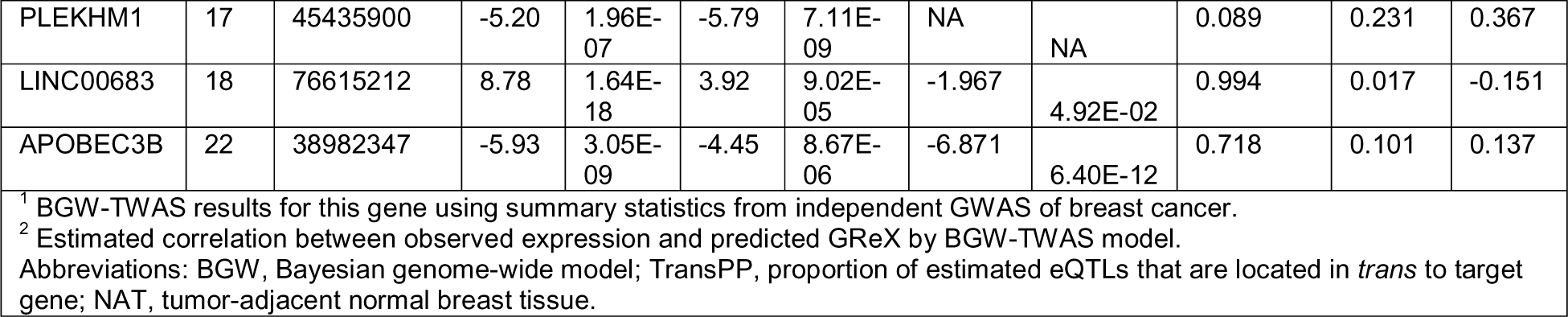
BGW-TWAS identified genes associated with risk of overall breast cancer in BCAC analysis that validated in Rashkin et al. GWAS analysis (31).

|  |  |  | BGW |  |  |  | sPrediXcan |  |  |  |  |
| --- | --- | --- | --- | --- | --- | --- | --- | --- | --- | --- | --- |
|  |  |  | BCAC GWAS |  | Validation GWAS<sup>1</sup> |  | BCAC GWAS |  |  | TCGA Spearman's rho<sup>2</sup> |  |
| Gene | Chr | Pos | Z | p | Z | p | Z | p | TransPP | Tumor | NAT |
| ACAP3 | 1 | 1292376 | 12.23 | 2.25E-34 | 5.80 | 6.68E-09 | -0.124 | 9.01E-01 | 0.996 | 0.031 | -0.048 |
| BLACAT1 | 1 | 205434886 | 9.14 | 6.46E-20 | 3.65 | 2.66E-04 | 0.523 | 6.01E-01 | 0.999 | -0.081 | 0.135 |
| ITGA10 | 1 | 145891208 | 6.91 | 4.71E-12 | 3.58 | 3.43E-04 | NA | NA | 0.995 | 0.064 | -0.13 |
| TMEM50A | 1 | 25338214 | -5.91 | 3.46E-09 | -4.66 | 3.17E-06 | -0.603 | 5.47E-01 | 0.992 | 0.031 | -0.011 |
| AC093110.3 | 2 | 54662556 | -7.98 | 1.44E-15 | -3.95 | 7.79E-05 | NA | NA | 0.882 | NA | NA |
| ALS2CR12 | 2 | 201288271 | 6.15 | 7.66E-10 | 5.06 | 4.10E-07 | 6.074 | 1.25E-09 | 0.705 | NA | NA |
| CYBRD1 | 2 | 171522247 | -5.77 | 7.83E-09 | -4.27 | 1.94E-05 | -5.165 | 2.41E-07 | 0.3 | 0.058 | 0.538 |
| KLF7-IT1 | 2 | 207120884 | 5.54 | 2.98E-08 | 4.39 | 1.14E-05 | NA | NA | 0.92 | 0.002 | 0.105 |
| SLC4A7 | 3 | 27372721 | 10.44 | 1.69E-25 | 4.94 | 7.79E-07 | 1.914 | 5.56E-02 | 0.838 | 0.116 | 0.112 |
| GLRA3 | 4 | 174636914 | -8.09 | 5.99E-16 | -3.96 | 7.41E-05 | NA | NA | 0.362 | 0.043 | -0.051 |
| RPS23 | 5 | 82273358 | 6.81 | 9.83E-12 | 5.15 | 2.65E-07 | 6.255 | 3.98E-10 | 0.042 | 0.395 | 0.553 |
| ATP6AP1L | 5 | 82279462 | -6.80 | 1.08E-11 | -5.14 | 2.81E-07 | -5.586 | 2.32E-08 | 0.288 | 0.296 | 0.533 |
| MRPS30 | 5 | 44808925 | 6.17 | 6.88E-10 | 3.74 | 1.82E-04 | 9.24 | 2.47E-20 | 0.804 | 0.1 | 0.066 |
| ATG10 | 5 | 81972025 | -6.14 | 8.04E-10 | -4.48 | 7.57E-06 | -4.835 | 1.33E-06 | 0.383 | 0.474 | 0.608 |
| NUDT1 | 7 | 2242222 | 6.18 | 6.29E-10 | 6.06 | 1.38E-09 | 0.072 | 9.43E-01 | 0.918 | -0.065 | -0.028 |
| NPM2 | 8 | 22024125 | -9.48 | 2.50E-21 | -5.32 | 1.02E-07 | -0.276 | 7.82E-01 | 0.999 | 0.036 | -0.176 |
| EFR3A | 8 | 131904088 | -7.51 | 6.16E-14 | -4.63 | 3.64E-06 | -0.555 | 5.79E-01 | 0.988 | -0.014 | 0.085 |
| RP11-723D22.3 | 8 | 38844866 | 5.33 | 9.93E-08 | 3.53 | 4.17E-04 | NA | NA | 0.995 | NA | NA |
| IL2RA | 10 | 6010689 | -6.73 | 1.66E-11 | -3.71 | 2.08E-04 | -1.084 | 2.78E-01 | 0.93 | -0.009 | -0.169 |
| RP11-165A20.3 | 10 | 24953241 | 6.66 | 2.78E-11 | 4.70 | 2.66E-06 | NA | NA | 0.999 | NA | NA |
| PIDD1 | 11 | 799191 | 7.33 | 2.25E-13 | 3.79 | 1.48E-04 | 7.249 | 4.19E-13 | 0.25 | 0.083 | 0.189 |
| CCDC91 | 12 | 28133249 | -6.78 | 1.22E-11 | -3.64 | 2.74E-04 | -4.391 | 1.13E-05 | 0.641 | 0.035 | 0.178 |
| RCCD1 | 15 | 90955796 | -8.58 | 9.50E-18 | -4.85 | 1.23E-06 | -8.208 | 2.25E-16 | 0.023 | 0.111 | 0.345 |
| RP11-467L19.16 | 15 | 20642799 | -5.87 | 4.37E-09 | -4.06 | 4.85E-05 | NA | NA | 1 | NA | NA |
| LINC02210 | 17 | 45620328 | -5.99 | 2.12E-09 | -6.00 | 1.98E-09 | NA | NA | 0.501 | 0.322 | 0.686 |
| RP11-259G18.1 | 17 | 46267037 | -5.93 | 2.97E-09 | -5.90 | 3.62E-09 | NA | NA | 0.002 | NA | NA |
| KANSL1-AS1 | 17 | 46193576 | -5.50 | 3.82E-08 | -5.99 | 2.10E-09 | NA | NA | 0.508 | 0.583 | 0.699 |
| CTD-3157E16.1 | 17 | 15787787 | -5.42 | 6.11E-08 | -4.02 | 5.71E-05 | NA | NA | 0.799 | NA | NA |
| PLEKHM1 | 17 | 45435900 | -5.20 | 1.96E-07 | -5.79 | 7.11E-09 | NA | NA | 0.089 | 0.231 | 0.367 |
| LINC00683 | 18 | 76615212 | 8.78 | 1.64E-18 | 3.92 | 9.02E-05 | -1.967 | 4.92E-02 | 0.994 | 0.017 | -0.151 |
| APOBEC3B | 22 | 38982347 | -5.93 | 3.05E-09 | -4.45 | 8.67E-06 | -6.871 | 6.40E-12 | 0.718 | 0.101 | 0.137 |

<sup>1</sup> BGW-TWAS results for this gene using summary statistics from independent GWAS of breast cancer.
<sup>2</sup> Estimated correlation between observed expression and predicted GReX by BGW-TWAS model.
Abbreviations: BGW, Bayesian genome-wide model; TransPP, proportion of estimated eQTLs that are located in *trans* to target gene; NAT, tumor-adjacent normal breast tissue.

For this overall analysis, the most significant gene identified by BGW-TWAS was *ACAP3* on chromosome 1 (BCAC p = 2.3 × 10^−34^). We do note that this gene was found to be similarly associated with risk of luminal-A-like breast cancer (BCAC p = 3.4 × 10^−34^). The BVSR genome-wide model contained 18 *cis*-SNPs and 1,052 *trans*-SNPs. As the upper plot of Figure 2 and Supplemental Figure 10 illustrate, the SNPs with highest posterior probability of being eQTLs and largest expected eQTL weights lie on chromosomes 10 and 5, and several of these SNPs on chromosome 10 additionally have highly significant GWAS p-values (Figure 2, bottom plot). Given that the association of this gene is predominantly driven by *trans*-eQTL effects, it was not identified by sPrediXcan in either the overall analysis or the luminal-A-like analysis. The SNP with the highest estimated probability of being an eQTL for *ACAP3* was rs1268974 on chromosome 10, and we further observe that many of the SNPs most likely to be eQTLs on chromosome 10 are intron variants for *ZMIZ1*. Notably, rs1268974 (PP = 0.1153) has been associated with breast cancer in European ancestry^2^. Another top predictor at this locus is rs704010 (PP = 0.0394), which has been associated with overall breast cancer in European ancestry^2^ and Han Chinese^51^ populations, as well as overall and ER-/PR-breast cancer in an African American cohort^52^.

**Figure 2.**
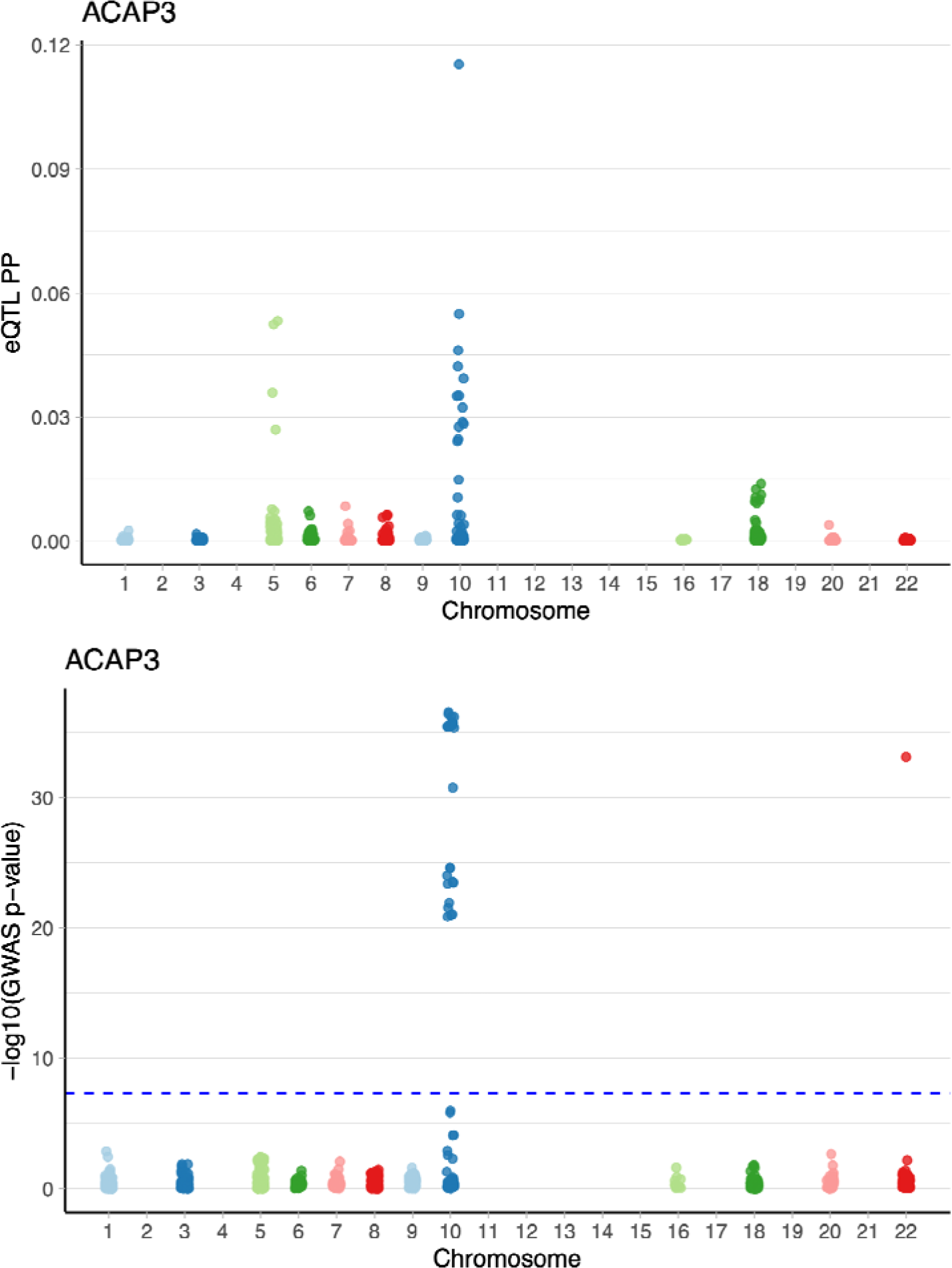
Estimated posterior probability (PP) of non-zero eQTL effects sizes from BGW-TWAS -selected SNPs for ACAP3 on chromosome 1 in breast tissue (top), and negative logarithm of the overall breast cancer GWAS p-values for these selected SNPs (bottom). Blue dotted line indicates genome-wide significance threshold for GWAS (5 × 10^−8^).

For our overall breast cancer analysis, several of the other top BGW-TWAS genes for this phenotype are supported by the literature. The majority of genes listed in Table 1 lie within 1Mb of one or more curated breast-cancer associated variants^53^. However, we consider 10 of these genes to represent novel candidate risk loci, as they do not lie near these variants or even a set of candidate susceptibility loci for ovarian cancer^8^. The expression models fitted for these genes (*ACAP3, CTD-3157E16.1, EFR3A, KLF7-IT1, LINC00683, NPM2, NUDT1, RP11-467L19.16, RP11-723D22.3, TMEM50A*) all show predominantly *trans*-SNP effects on regulation. The proportion of estimated eQTLs from *trans* regions exceeds 0.79 for each gene, and none were identified by sPrediXcan.

Beyond *ACAP3*, we identified 60 additional genes in the subtype-specific analysis of luminal A-like cancer using BGW-TWAS with BCAC data. Of these, 53 were also among those identified above for overall breast cancer risk. It is important to note that the independent GWAS summary data that we used for validation analyses of our BCAC-derived findings did not distinguish cases by cancer subtype. As such, we are less powered to validate certain subtype-specific associations, particularly for those of rarer cancer types. However, given the high prevalence of luminal A-like cancers globally, we assume the GWAS phenotype of the validation data (overall cancer) to be a reasonably proxy for the luminal-A-like phenotype. In Table 2, we present the 18 luminal-A-like genes that further validated when we applied our BGW-TWAS models to the Rashkin et al. data (p < 0.05/101). Of these 18 genes, *ACAP3, EFR3A, NPM2, NUDT1,* and *RP11-723D22.3* are also not in regions of candidate breast cancer susceptibility loci^53^. The full list of all TWAS genes identified for luminal A-like cancer by BGW-TWAS are provided in Supplemental Table 3.

**Table 2.**
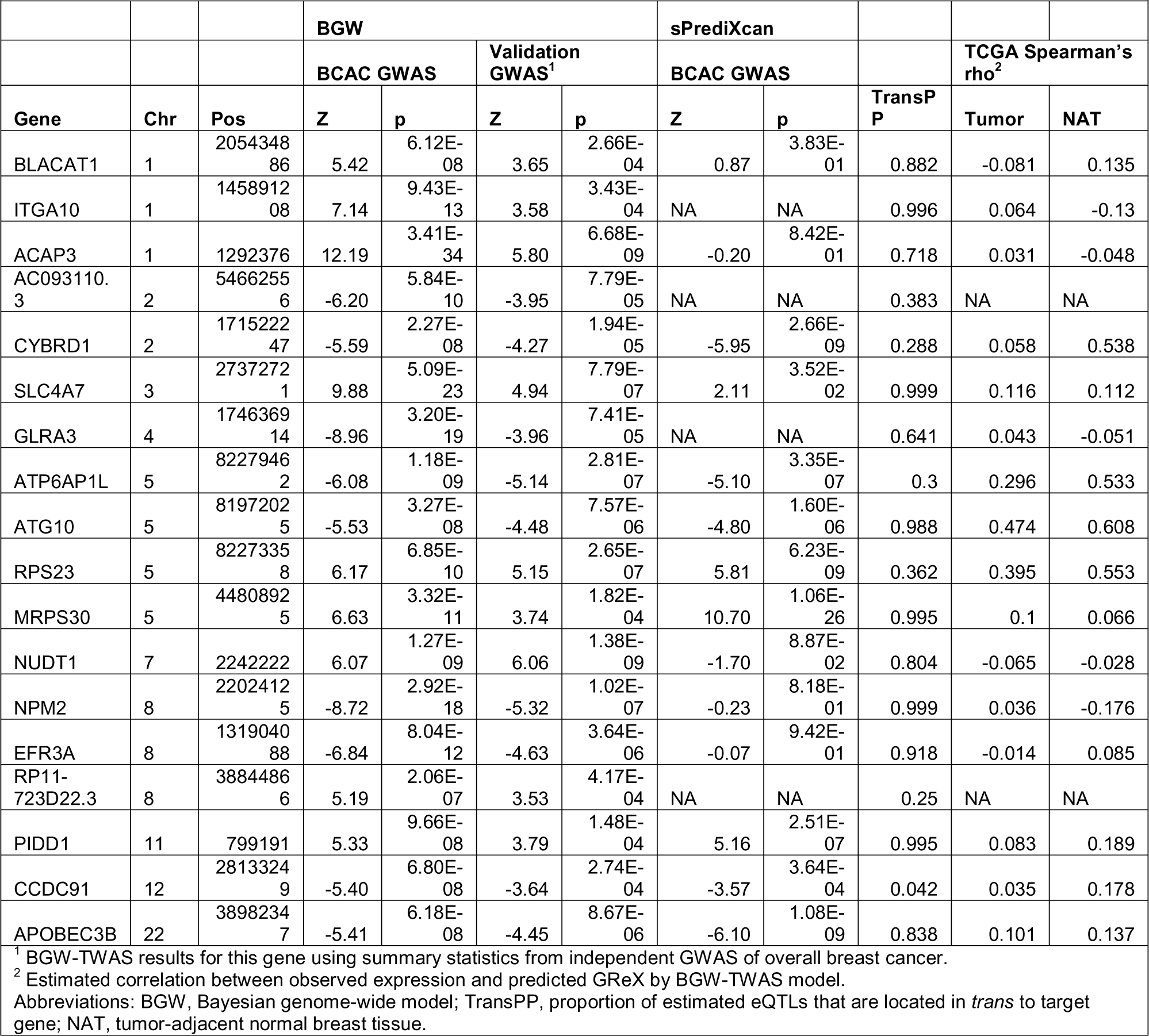
BGW-TWAS identified genes associated with risk of luminal-A-like breast cancer in BCAC analysis that validated in Rashkin et al. GWAS analysis (18).

Results from the TWAS of all other breast cancer subtypes (luminal B-like, luminal B/HER2-negative-like, HER2-enriched-like, and triple-negative) are shown in Table 3. We do not restrict this table to those genes that additionally had a significant association using the validation data since, here, we do not expect an independent GWAS of overall breast cancer to be an ideal validation study for these subtype-specific findings. However, the corresponding effect size estimates and p-values from application of BGW-TWAS to the validation summary data are provided in Table 3. The subtype-specific risk genes we identified also reflect prior candidate loci. For example, previous work has suggested that *BLACAT1* has a role in breast cancer metastasis^54^. Expression of *RCCD1* on chromosome 15 has also been shown to be associated with breast cancer risk in a large trans-ethnic TWAS^55^. However, *BOD1L1* (luminal B-like), *NPM2* (luminal B/HER2-negative-like), *RP11-474P2.6* (HER2-enriched-like) and *RPS18* (triple-negative) do not fall within 1Mb of curated candidate breast cancer risk loci^53^. For a global comparison of findings across all our analyses in breast cancer, we provide a correlation plot of BGW-TWAS Z-scores across all genes between subtypes in Supplemental Figure 11.

**Table 3.** BGW-TWAS identified genes associated with specific risk of other breast cancer subtypes in BCAC analysis with corresponding PrediXcan and validation results.

| Subtype | Gene | Chr | Pos | TransPP | BGW p<br>BCAC<br>GWAS | BGW p<br>Validation<br>GWAS<sup>†</sup> | PrediXcan<br>p<br>BCAC<br>GWAS |
| --- | --- | --- | --- | --- | --- | --- | --- |
| Luminal B-like | BOD1L1 | 4 | 13568738 | 0.217 | 2.70E-09 | 0.354 | 2.09E-09 |
|  | RCCD1 | 15 | 90955796 | 0.023 | 8.33E-08 | 1.23E-06 | 2.48E-07 |
| Luminal B/HER2-negative-like | NPM2 | 8 | 22024125 | 0.999 | 3.81E-10 | 1.02E-07 | 0.590 |
|  | RCCD1 | 15 | 90955796 | 0.023 | 1.60E-12 | 1.23E-06 | 1.86E-12 |
| HER2-enriched like | BLACAT1 | 1 | 205434886 | 0.999 | 2.64E-08 | 0.000266 | 0.365 |
|  | RP11-474P2.6 | 12 | 46388856 | 0.998 | 1.09E-07 | 0.751208 | 0.641 |
| Triple-negative | BLACAT1 | 1 | 205434886 | 0.999 | 4.45E-10 | 0.000266 | 0.613 |
|  | RPS18 | 6 | 33272048 | 0.391 | 5.85E-10 | 0.773831 | 1.43E-12 |
|  | ANKLE1 | 19 | 17281645 | 0.378 | 1.28E-41 | 0.004143 | 4.21E-30 |
|  | OCEL1 | 19 | 17226204 | 0.190 | 2.15E-14 | 0.004081 | 9.71E-12 |
|  | ABHD8 | 19 | 17292131 | 0.001 | 4.73E-09 | 0.741252 | 1.31E-20 |

<sup>†</sup> BGW-TWAS results for this gene using summary statistics from independent GWAS of overall breast cancer.
Abbreviations: BGW, Bayesian genome-wide model; TransPP, proportion of estimated eQTLs that are located in *trans* to target gene.

For breast cancer genes, we additionally used the GTEx-derived models of genetically regulated gene expression to predict gene expression levels in tumor tissue samples of breast cancer patients from TCGA, as well as normal tumor-adjacent breast tissue samples. This analysis was performed to gauge how well the models trained in GTEx among individuals without breast cancer accurately predict transcription in individuals with breast cancer. Tables 1-3 provide an estimate of the correlation between imputed gene expression in TCGA samples that used the GTEx-derived BGW-TWAS models and observed expression levels in both tumor and NAT breast tissue. *RPS23, ATP6AP1L, ATG10, RCCD1, LINC02210*, and *KANSL1-AS1* showed nominally significant correlation between imputed and observed expression levels in both NAT and tumor TCGA samples (p < 0.05), and the estimated Spearman correlation coefficient was positive between the two vectors. The *SLC4A7* model validated in tumor tissue, but not NAT. We also saw validation of the models in tumor tissue only for *THBS3*, significant in the analysis of luminal A-like cancer, and *RPS18*, identified for triple-negative-like breast cancer. Further, the BGW models of *MAN2C1* (luminal A-like cancer) and *ABHD8* (triple negative) showed significant correlation between predicted and observed expression in NAT tissue only. Three other significant genes identified in the subtype-specific TWAS validated in both tissue types in TCGA: *ATE1-AS1* (luminal A-like), *RCCD1* (luminal B-like, luminal B/HER2-negative-like), and *OCEL1* (triple-negative).

### 3.3 Ovarian cancer TWAS

We performed a total of 135,474 tests using BGW-TWAS across the six ovarian cancer phenotypes. This corresponds to a Bonferroni threshold of 3.69 ×10^−7^ for transcriptome-wide significance. We obtained sPrediXcan results for 13,109 genes for each of the six ovarian cancer phenotypes, corresponding to a total of 78,654 tests and a Bonferroni threshold of 6.36×10^−7^. We present genome-wide findings for the non-mucinous ovarian cancer and the five main subtypes of ovarian cancer using BGW-TWAS and OCAC-derived GWAS summary statistics in Supplemental Figures 12-18. The BGW-TWAS p-values do show evidence of some inflation for the non-mucinous and high grade serous phenotypes. However, as with the breast cancer results, this is corrected when considering the GWAS sample size used in construction of the test statistics (λ = 1.06, λ_1000_ = 1.002 for NMOC).

Eight unique significant genes were identified by BGW-TWAS when applied to the summary statistic data on risk of NMOC and HGSOC (Table 4). No genes meeting the multi-trait adjusted Bonferroni threshold were identified for LGSOC, EOC, MOC, or CCOC. A correlation plot of BGW-TWAS Z-scores across all genes between ovarian cancer subtypes is included in Supplemental Figure 19. All eight significant genes had model training R^2^ > 0.1 in ovarian tissue. Of the eight genes shown, sPrediXcan fit models for *ANKLE1* and *CCDC106*. The most significant gene identified by BGW-TWAS across all analyses was *ANKLE1* at 19p13 for HGSOC (BCAC p = 4.4×10^−21^), but it was also identified in the non-mucinous analysis (BCAC p = 8.25×10^−13^). *ANKLE1* is a well-established candidate susceptibility locus for both breast cancer and ovarian cancer^31,56,57^. In the sPrediXcan model of *ANKLE1*, three SNPs were used to model expression, one of which (rs67412075) was also included in the BGW-TWAS model for *ANKLE1* in ovarian tissue (PP = 0.0169). Although 389 out of 448 selected SNPs were located in *trans* to *ANKLE1*, all SNPs driving the association (with highest eQTL PP) are *cis*-SNPs located on chromosome 19, and thus it is not surprising that this gene was similarly identified by sPrediXcan in both phenotypes.

**Table 4.**
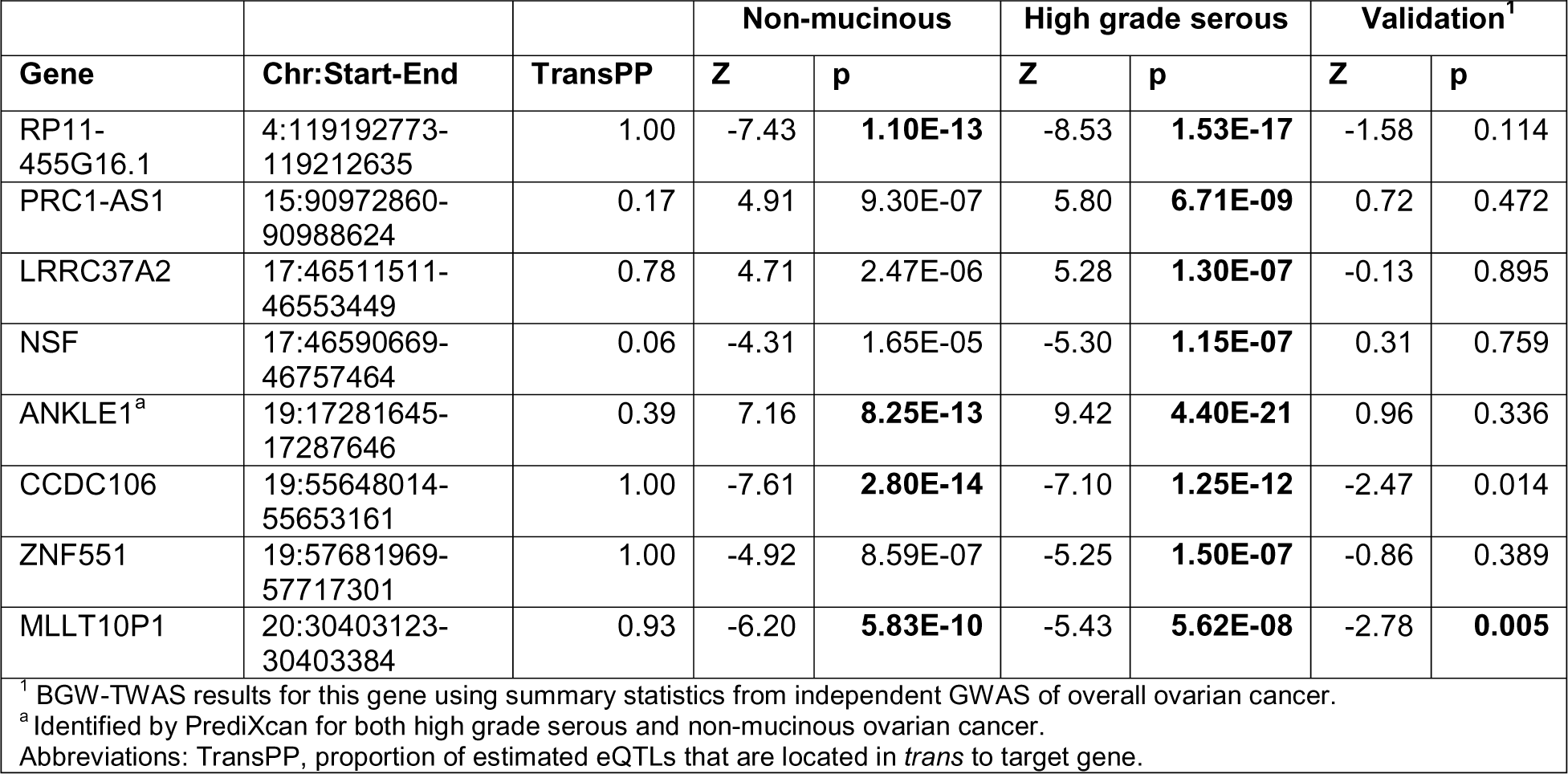
Significant genes identified by BGW-TWAS for non-mucinous and high grade serous ovarian cancer in OCAC analyses with validation p-value for overall ovarian cancer using Rashkin et al. GWAS summary statistics.

In contrast, another highly significant gene from the BGW-TWAS analysis of both NMOC and HGSOC that appears to be largely driven by *trans*-eQTL effects is *CCDC106* at 19q13. The BVSR genome-wide model had 21 *cis*-SNPs and 2016 *trans*-SNPs. As we can see in the top plot of Figure 3 and Supplemental Figure 20, the SNPs with highest PP of being eQTLs and largest expected eQTL weights are located in *trans* on chromosomes 17, 2, and 15. The bottom figure of Figure 3, which shows the corresponding OCAC GWAS p-values for these SNPs in the non-mucinous analysis, indicates that a subset of SNPs on chromosome 17 furthermore have the most significant GWAS associations for this phenotype. Therefore, this gene has not been implicated in previous *cis*-only TWAS of ovarian cancer. *CCDC106* was not identified as significant by sPrediXcan for either HGSOC or NMOC (p > 0.2), as the association appears to be driven by distal GWAS-significant loci on chromosome 17 correlated with *CCDC106* expression. The top SNP by eQTL PP is rs1979858 on chromosome 15, an intergenic variant for *ARRDC4*. The second top SNP is rs9898988 on chromosome 17, an intron variant for *SKAP1*. Intron variants of *SKAP1* have previously been associated with overall epithelial ovarian cancer^8,58,59^ and HGSOC^8^.

**Figure 3.**
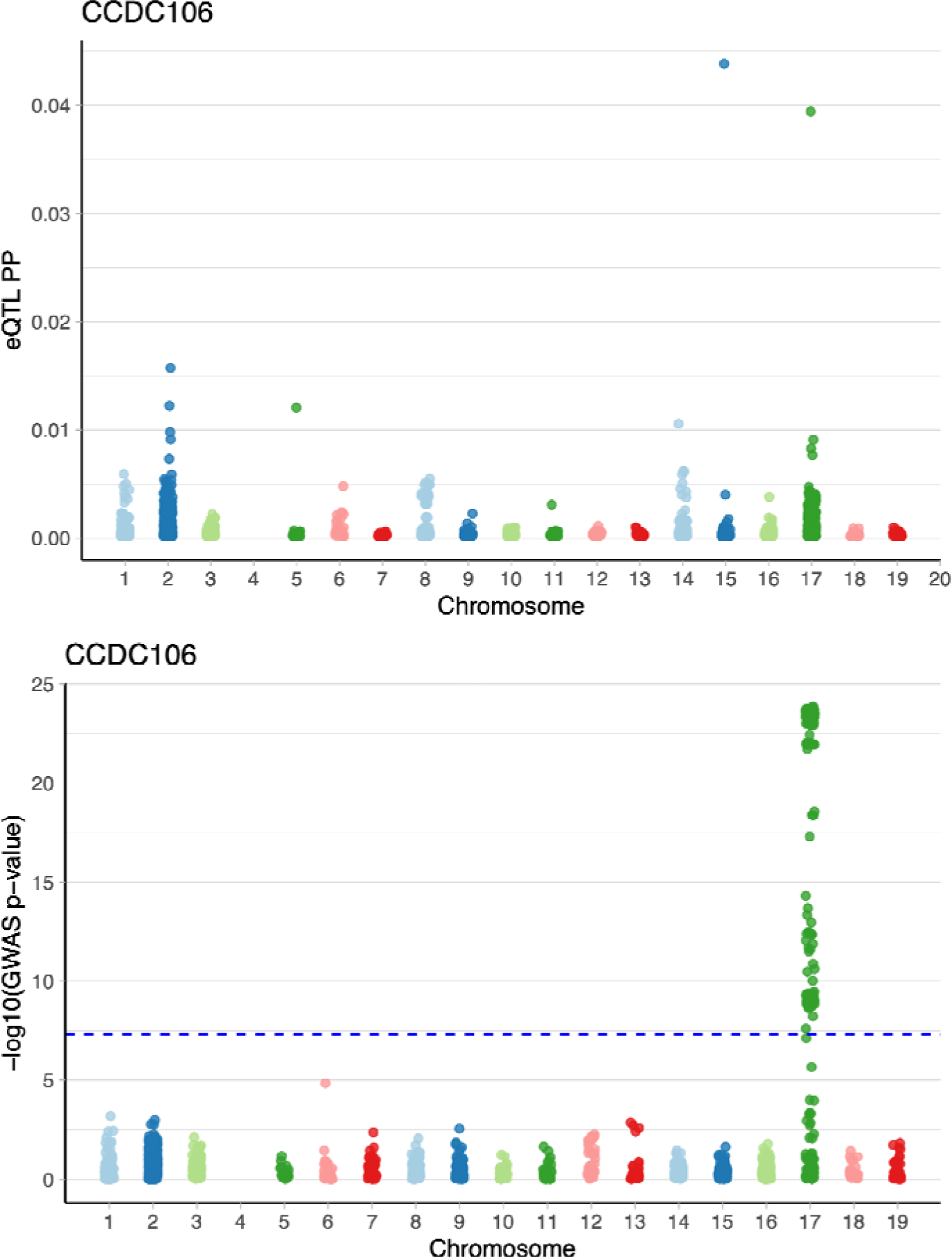
Estimated posterior probability (PP) of non-zero eQTL effects sizes from BGW-TWAS -selected SNPs for CCDC106 on chromosome 19 in ovarian tissue (top), and negative logarithm of the non-mucinous ovarian cancer GWAS p-values for these selected SNPs (bottom). Blue dotted line indicates genome-wide significance threshold for GWAS (5 × 10^−8^).

Other top genes identified by BGW-TWAS have been implicated by previous work. For example, *PRC1-AS1* was identified for HGSOC and is a candidate risk locus for breast cancer previously identified by GWAS in Europeans and East Asians^60,61^ and also identified by a previous *cis*-only TWAS of ovarian cancer^23^. Additionally, for the HGSOC analysis, our genome-wide model implicates *NSF*, which has documented GWAS and TWAS associations with risk of ovarian cancer^23,62^. For *R11-455G15.1*, the top SNP by eQTL PP was rs10738466 on chromosome 9. This SNP is 32,525 bp to *BNC2*, a locus already implicated in ovarian cancer^59,63–65^.

However, of the eight ovarian cancer genes identified across phenotypes, *LRRC37A2, CCDC106, ZNF551*, and *MLLT10P1* do not lie within 1 Mb of reported ovarian susceptibility loci^8^. *CCDC106* and *MLLT10P1* are additionally not within regions of curated breast cancer susceptibility loci^53^. For *MLLT10P1*, the top SNP most likely to be an eQTL was rs2229304 on chromosome 17. This is a missense variant for *HOXB23. MLLT10P1* was additionally the only gene to validate when BGW-TWAS was applied to independent GWAS summary statistics for overall ovarian cancer risk by Rashkin et al. (p < 0.05/8). We note that failure of all other genes to replicate is likely due in part to the limited sample size for ovarian cancer cases in this validation GWAS itself (1,006 UKB, 253 GERA). These counts are considerably smaller than the corresponding sample size of breast cancer cases used (13,903 UKB, 3,978 GERA) and overall controls used (189,855 UKB, 29,801 GERA). Although only one gene achieved significance in this validation, six out of the eight genes originally identified by BGW-TWAS with OCAC summary data showed the same estimated direction of effect in the follow-up validation analysis (Table 4). For the two genes with differing effect directions between the OCAC and Rashkin et al. analyses, we emphasize that the latter Z-scores were close to zero (p > 0.7).

## 4 DISCUSSION

In this work, we conducted the first TWAS of breast cancer and ovarian cancer that uses not only *cis*-SNPs, but both intra- and inter-chromosomal *trans*-SNPs, to model genetically regulated transcription. This genome-wide modeling approach, BGW-TWAS, stems from the growing catalog of *trans*-eQTL effects identified across a wide range of tissue types^26,66–68^. We applied this method to train gene expression models in GTEx mammary and ovarian tissues and tested for association with risk of breast and non-mucinous ovarian cancer using summary GWAS data from recent large-scale meta-analyses. We further investigated how the landscape of identified risk genes for these diseases varied across cancer subtypes. We identified 101 significant genes across the overall and subtype-specific breast cancer analyses and 8 for the corresponding ovarian cancer analyses.

Although many of these loci fall nearby known GWAS risk variants or have been similarly implicated by other TWAS, several genes appear to be novel associations that are driven largely by *trans*-eQTL effects. ACAP3, EFR3A, NPM2, and NUDT1 were (1) identified in our TWAS for both luminal A-like breast cancer and overall breast cancer, (2) did not lie near curated sets of candidate susceptibility variants for either breast or ovarian cancer, and (3) further validated using an independent GWAS dataset. KLF7-IT1, LINC00683, and TMEM50A further met this criteria for overall breast cancer only. *ACAP3* is predicted to play a role in GTPase activator activity, but the gene’s possible role in tumorigenesis is unknown. *EFR3A* protein, however, has been implicated in oncogenic signaling and tumorigenic activity^69^. NUDT1 overexpression has been observed in several cancers, including breast^70,71^. *NPM2* is located near a well-studied tumor suppressor gene, *DOK2,* on 8p21.3, which is theorized to play a role in several cancers^72^. *NPM2* further showed association with risk of luminal B/HER2-negative-like breast cancer using both BCAC and validation GWAS data.

While less powered to detect novel genes associated with risk of ovarian cancer and its main subtypes due to the limited number of ovarian tissue samples available in GTEx, we did identify two genes that may warrant further investigation. *CCDC106* and *MLLT10P1* were strongly associated with both HGSOC and NMOC with *trans*-driven GReX. They are not located near sets of curated candidate breast cancer and ovarian cancer risk variants. However, recent work in mutant p53 ovarian cancer cells has shown that overexpression of *CCDC106* in particular leads to inhibition of p21 transcription and, ultimately, proliferation of the cancer cells^73^. *MLLT10P1* was the only risk gene to validate using independent GWAS data in our ovarian cancer analyses. While it is a pseudogene, and therefore the biological mechanisms behind this association are unclear, functional research on pseudogenes has indicated that they can indeed play a role in tumorigenesis and are dysregulated in many cancers^74^.

In this study, the abundance of non-trivial *trans*-SNP effects on gene expression that we observed in both mammary and ovarian tissues opened the door for identification of new potential risk genes and underscores the importance of including *trans*-SNPs in TWAS. However, we note there are several limitations to this work. Firstly, the samples used for training the genome-wide expression imputation models were limited in number, particularly for ovarian tissue (N=140). One possible consequence of such a modest training size is overfitting, as reflected by inflated R^2^ in the training samples. While BGW-TWAS is the first method of its kind to be computationally tractable enough to fit *trans*-eQTL models, its computational requirements do prevent cross-validation analysis during model training. Also, the lower prevalence of ovarian cancer relative to breast cancer makes identification of suitable validation GWAS datasets and transcriptomic panels in ovarian tissue of sufficient sample size difficult to obtain.

Furthermore, we note that while breast cancer is a disease predominantly occurring among females, the majority of the GTEx samples for which RNA sequencing data was available in breast tissue came from men (63%). While training gene expression models using only female samples is ideal, the subsequent drop in sample size would have negatively impacted model predication accuracy. We also limited our scope to autosomal genes only and did not consider the important role of sex-chromosome genes in sex-biased diseases like breast cancer. Our study also used data only from individuals of European ancestry for the expression model training, gene-level association tests, and validation analyses. However, there are considerable disparities in clinical outcomes of these cancers across racial groups. Research has also reported unique gene expression profiles across non-European ancestries in breast cancer tumors^75^, which motivates that the application of *trans*-eQTL TWAS to underrepresented populations.

Lastly, we note that the expression models of our *trans*-eQTL-driven TWAS genes were trained using normal breast and ovarian tissue from GTEx rather than tumor adjacent normal tissue or tissue with precursor lesions that are disease relevant. Our expression models showed little validation in tumor adjacent normal tissue in TCGA, the closest independent surrogate samples for the GTEx normal breast tissue used to train our models. This may be a result of sample size or due to altered regulatory effects of these SNPs in these samples caused by their proximity to tumors. Indeed, recent work comparing GTEx tissues and the corresponding TCGA NAT across cancer types suggests that the NAT transcriptome is not “normal” but represents an intermediate gene expression state between normal and tumor with multiple pathway-level perturbations differentiating NAT from GTEx^76^. Furthermore, germline control of somatic gene expression may be lost during the oncogenic transition from normal (GTEx) to NAT and/or tumor tissue (TCGA)^77^. Regardless, other approaches are needed to validate the high probability *trans*-eQTLs of our novel risk genes. One possible extension would be analysis of Hi-C or other chromatin conformation capture data to quantify the rate at which our novel target genes and their corresponding regions of high-evidence *trans*-eQTLs interact physically in the nucleus. The 3D interaction of these regions in both normal breast tissue samples and established breast cancer cell lines would provide valuable insight into the complex genome-wide genetic regulation that we observe.

## Acknowledgements

This work was supported by the National Institutes of Health [AG071170, CA211574, GM138313].

## Web Resources

The BCAC GWAS summary statistic data for risk of breast cancer phenotypes are publicly available and can be accessed at https://bcac.ccge.medschl.cam.ac.uk/bcacdata/oncoarray/oncoarray-and-combined-summary-result/gwas-summary-associations-breast-cancer-risk-2020/. The manuscript for the OCAC GWAS of ovarian cancer phenotypes is currently under review, and the GWAS summary data will be made available for download upon publication. The Rashkin et al. pan-cancer GWAS summary statistics used in validation analyses are publicly available and can be accessed at https://github.com/Wittelab/pancancer_pleiotropy. The MetaXcan suite of tools are available for download on Github at https://github.com/hakyimlab/MetaXcan.

