## Supplemental Figures for "*Cis*- and *trans*-eQTL TWAS of breast and ovarian cancer identify more than 100 risk associated genes in the BCAC and OCAC consortia"

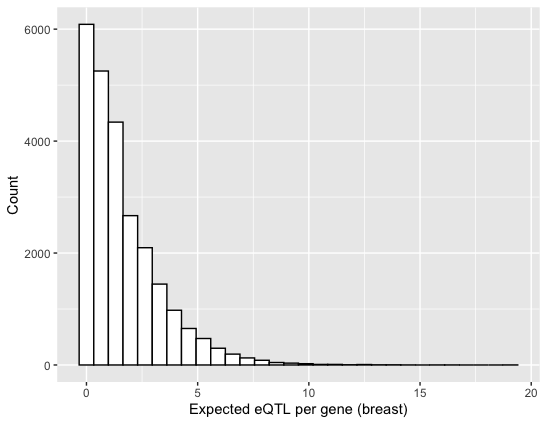

**Supplemental Figure 1**. Histogram of expected number of eQTL (cumulative posterior causal probability) per gene from BGW-TWAS imputation models in breast tissue.

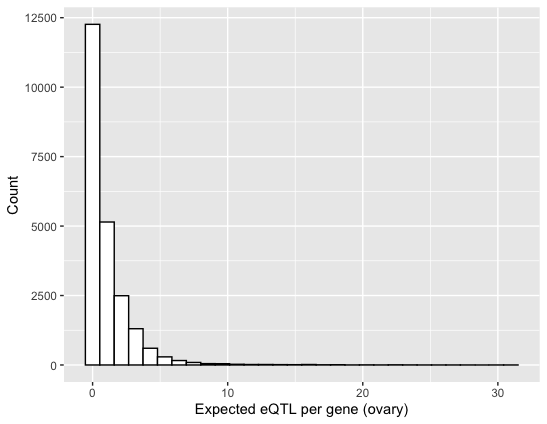

**Supplemental Figure 2**. Histogram of expected number of eQTL (cumulative posterior causal probability) per gene from BGW-TWAS imputation models in ovarian tissue.

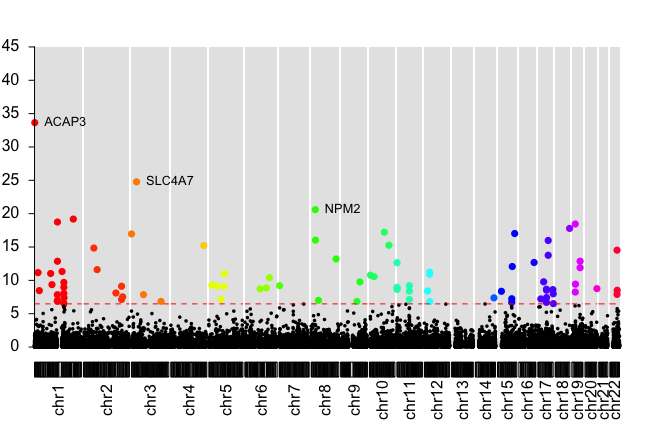

**Supplemental Figure 3**. Manhattan plot of BGW-TWAS results for overall breast cancer risk. The dashed line denotes the Bonferroni-adjusted transcriptome-wide significance threshold.

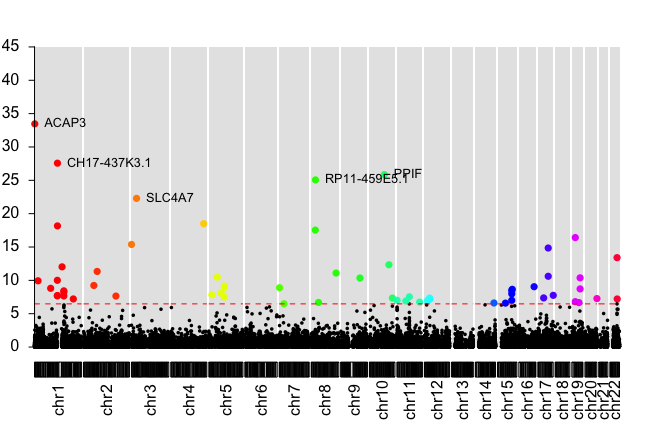

**Supplemental Figure 4.** Manhattan plot of BGW-TWAS results for luminal A-like breast cancer risk. The dashed line denotes the Bonferroni-adjusted transcriptome-wide significance threshold.

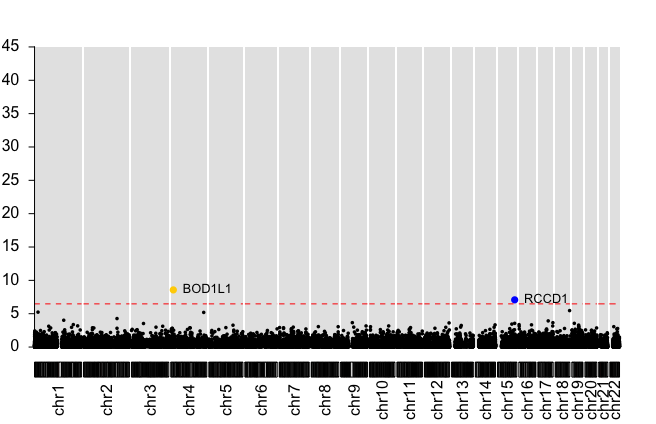

**Supplemental Figure 5**. Manhattan plot of BGW-TWAS results for luminal B-like breast cancer risk. The dashed line denotes the Bonferroni-adjusted transcriptome-wide significance threshold.

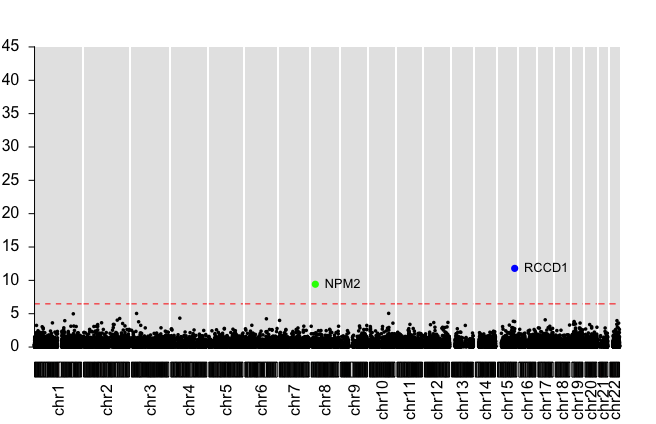

**Supplemental Figure 6**. Manhattan plot of BGW-TWAS results for luminal B/HER2 negative-like breast cancer risk. The dashed line denotes the Bonferroni-adjusted transcriptome-wide significance threshold.

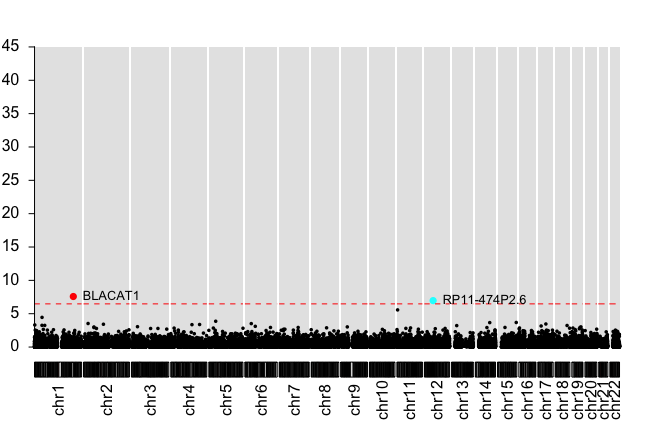

**Supplemental Figure 7**. Manhattan plot of BGW-TWAS results for HER2 enriched-like breast cancer risk. The dashed line denotes the Bonferroni-adjusted transcriptome-wide significance threshold.

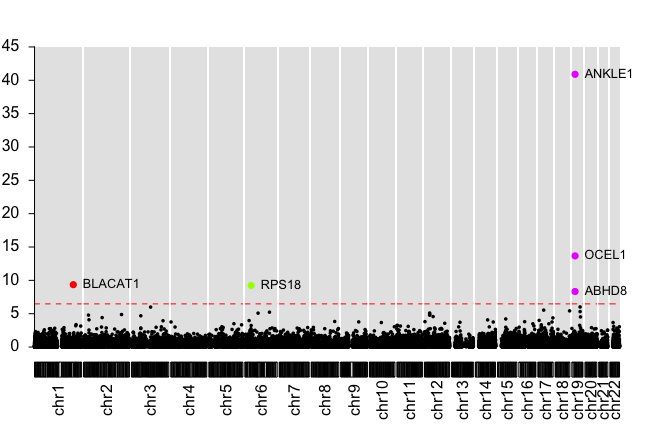

**Supplemental Figure 8**. Manhattan plot of BGW-TWAS results for triple negative-like breast cancer risk. The dashed line denotes the Bonferroni-adjusted transcriptome-wide significance threshold.

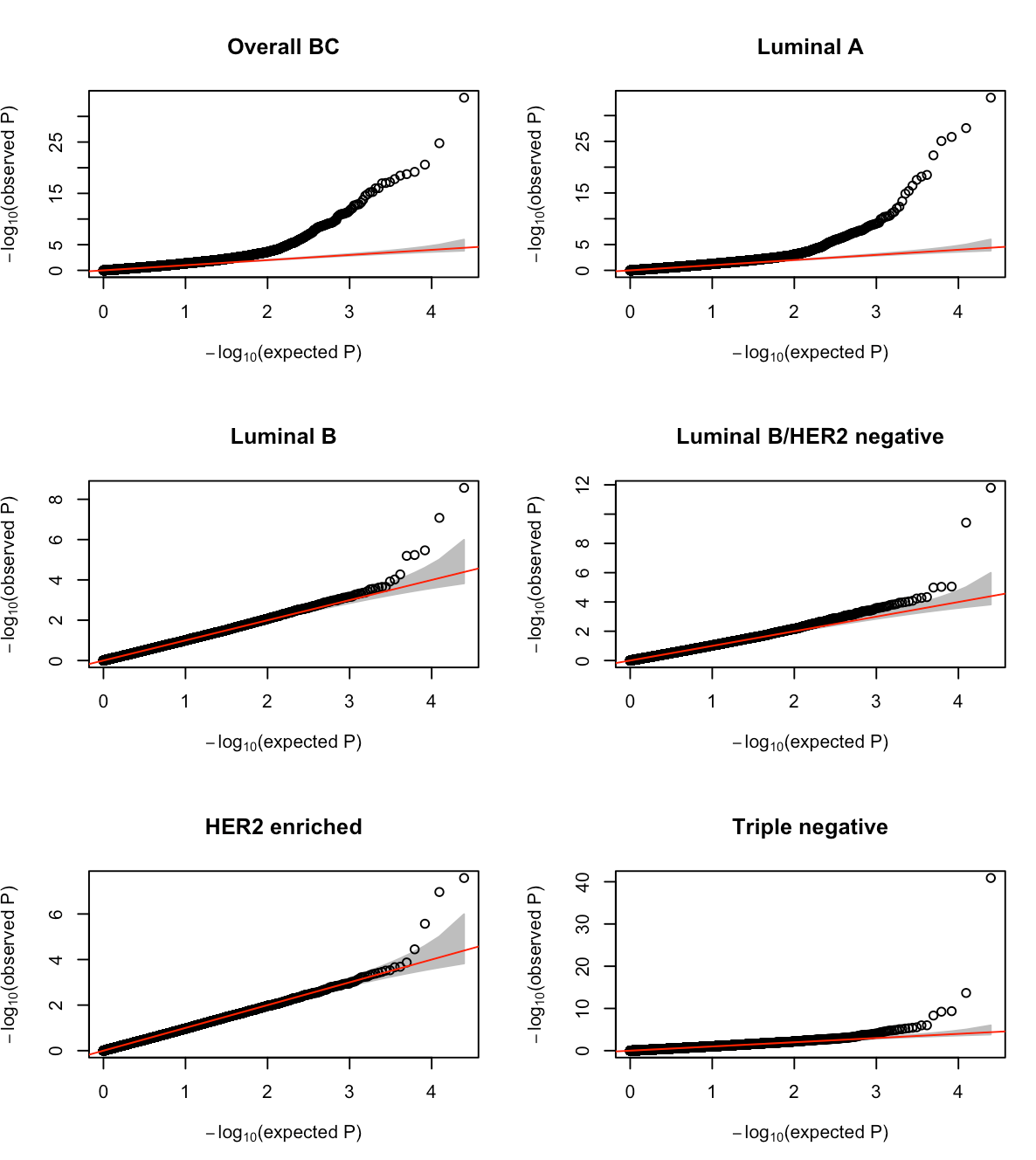

**Supplemental Figure 9.** Quantile-quantile plots of BGW-TWAS results for six breast cancer phenotypes using BCAC GWAS summary statistics. Abbreviations: BC, breast cancer.

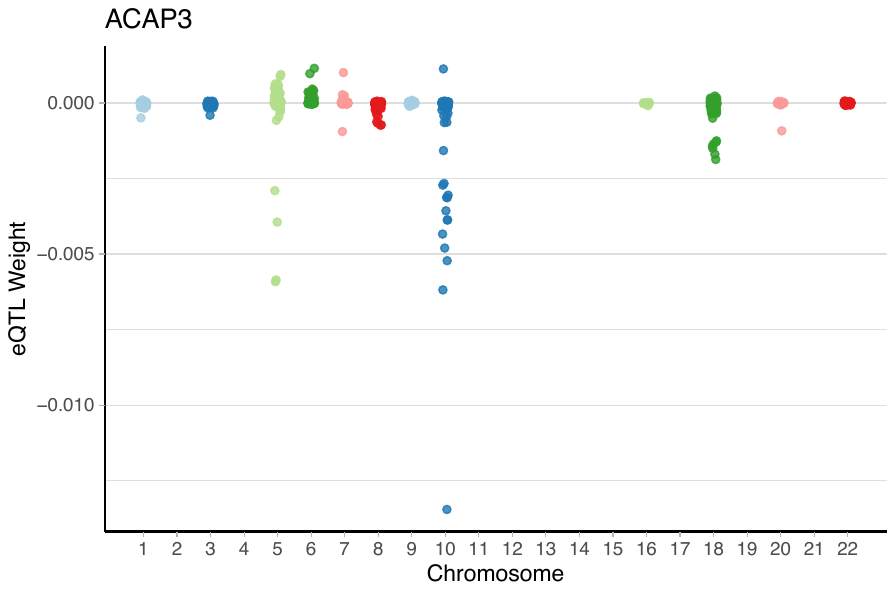
**Supplemental Figure 10**. Expected eQTL effect sizes estimated from the BVSR model for BGW-TWAS-selected SNPs for ACAP3 on chromosome 1 in breast tissue.

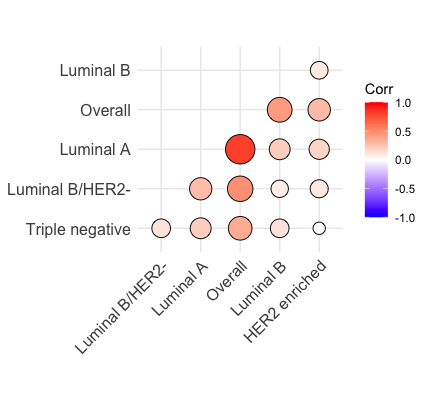

**Supplemental Figure 11.** Correlation plot of BGW-TWAS Z scores across six breast cancer phenotypes using BCAC GWAS summary statistics.

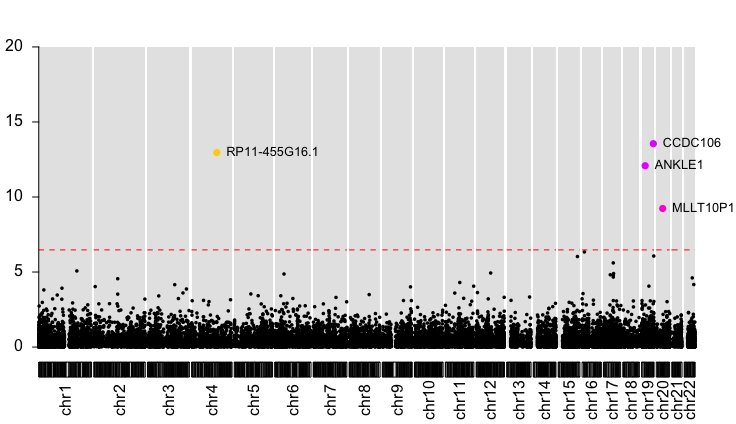

**Supplemental Figure** **12**. Manhattan plot of BGW-TWAS results for non-mucinous ovarian cancer risk. The dashed line denotes the Bonferroni-adjusted transcriptome-wide significance threshold.

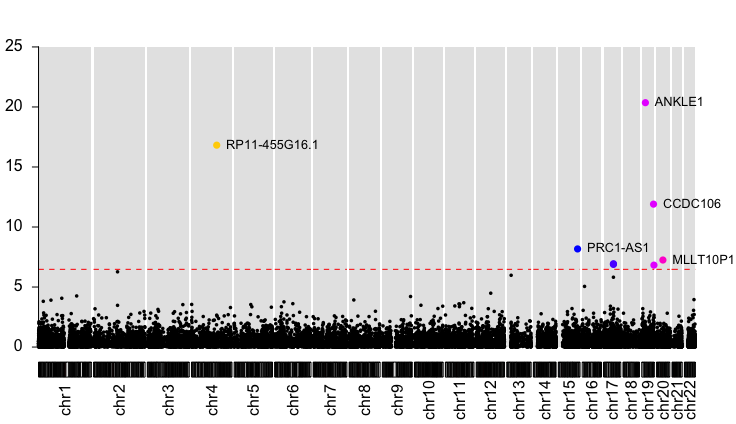

**Supplemental Figure 13**. Manhattan plot of BGW-TWAS results for high grade serous ovarian cancer risk. The dashed line denotes the Bonferroni-adjusted transcriptome-wide significance threshold.

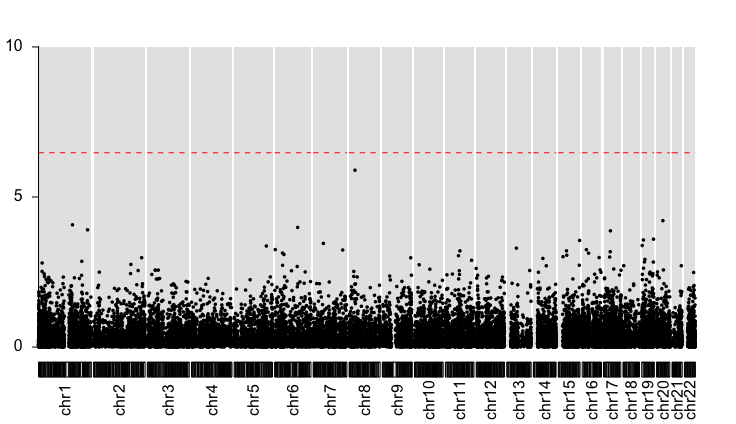

**Supplemental Figure 14**. Manhattan plot of BGW-TWAS results for low grade serous ovarian cancer risk. The dashed line denotes the Bonferroni-adjusted transcriptome-wide significance threshold.

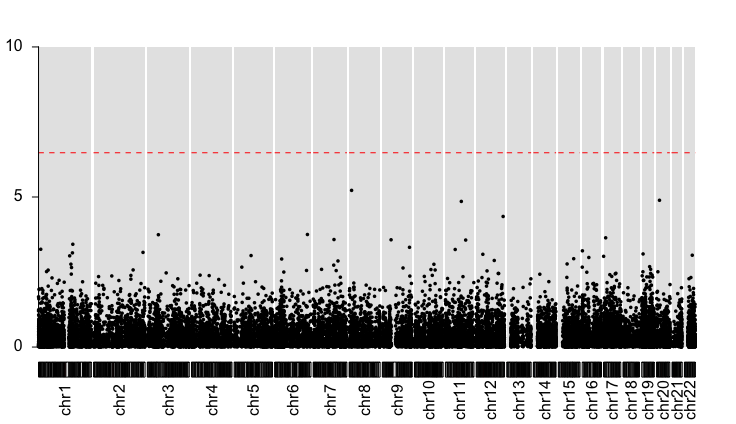

**Supplemental Figure 15**. Manhattan plot of BGW-TWAS results for mucinous ovarian cancer risk. The dashed line denotes the Bonferroni-adjusted transcriptome-wide significance threshold.

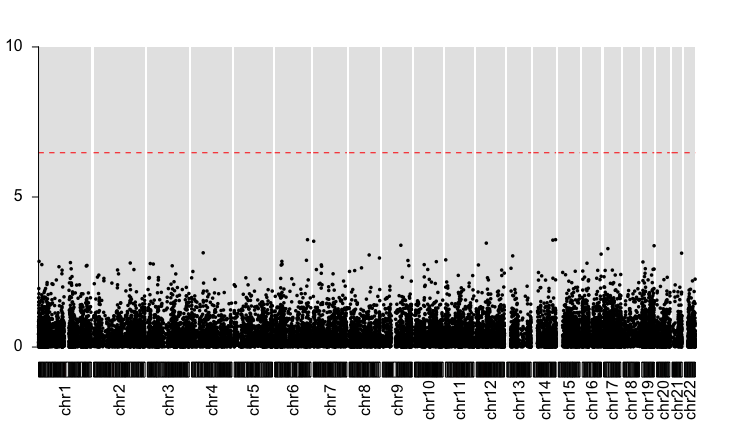

**Supplemental Figure 16**. Manhattan plot of BGW-TWAS results for endometrioid ovarian cancer risk. The dashed line denotes the Bonferroni-adjusted transcriptome-wide significance threshold.

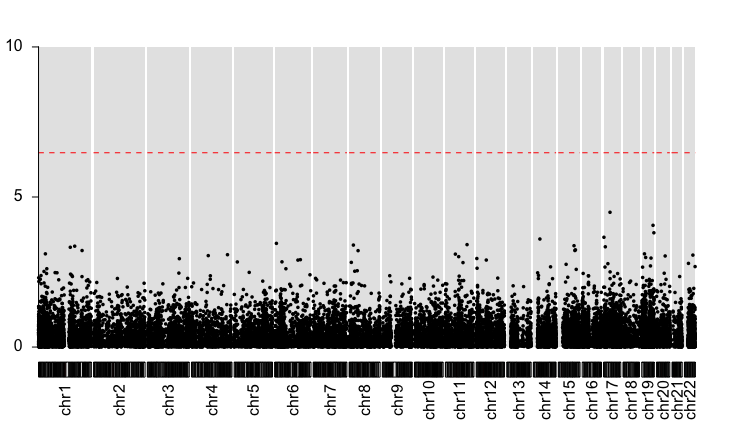

**Supplemental Figure 17**. Manhattan plot of BGW-TWAS results for clear cell ovarian cancer risk. The dashed line denotes the Bonferroni-adjusted transcriptome-wide significance threshold.

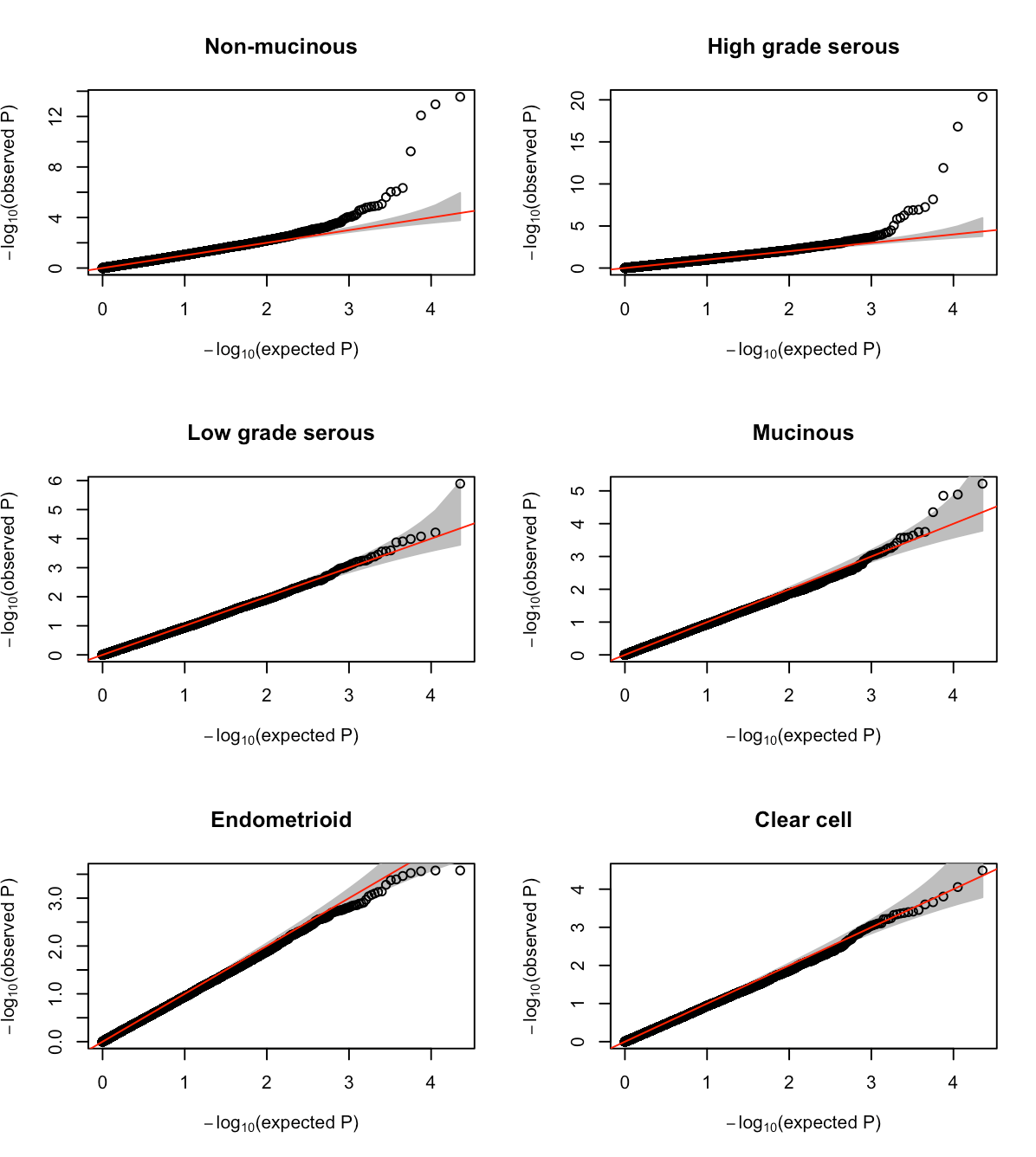

**Supplemental Figure 18.** Quantile-quantile plots of BGW-TWAS results for six ovarian cancer phenotypes using OCAC GWAS summary statistics.

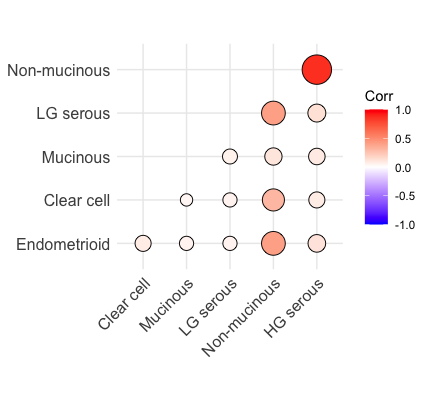

**Supplemental Figure 19**. Correlation plot of BGW-TWAS Z scores across six ovarian cancer phenotypes using OCAC GWAS summary statistics.

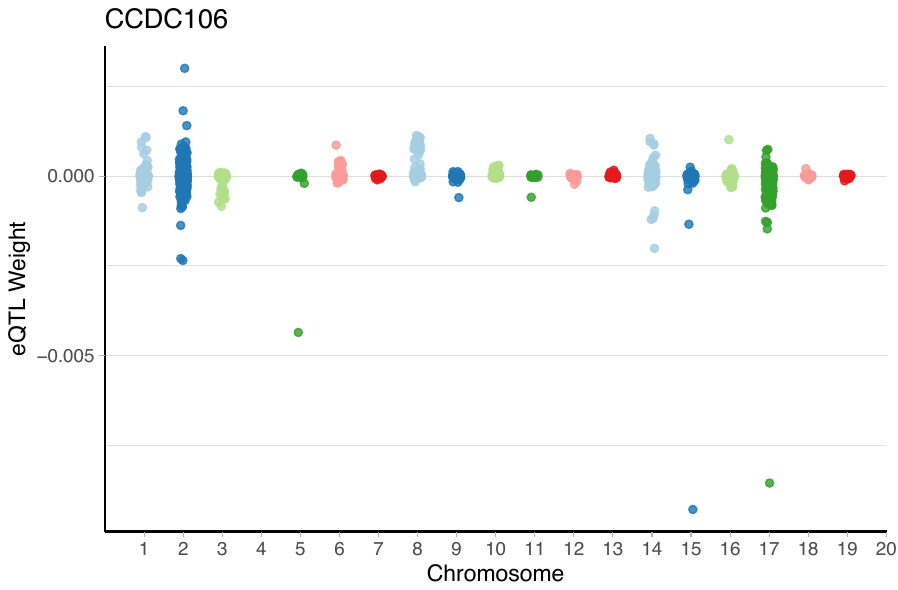

**Supplemental Figure 20**. Expected eQTL effect sizes estimated from BVSR model for BGW-TWAS-selected SNPs for CCDC106 on chromosome 19 in ovarian tissue.
